# Non-uniform structural development across human thalamus aligns with risk zones for schizophrenia in adulthood

**DOI:** 10.64898/2026.06.29.735354

**Authors:** Omar Singleton, Jesse Gomez

## Abstract

With dense axonal connectivity to every region of cortex, the thalamus plays a central role in the nervous system from sensory processing to cognitive functions. Yet how tissue maturation of the thalamus unfolds during childhood and contributes to typical or atypical development is not clear. Through several large datasets, we provide here a thalamic portrait of fine-scale structural development whose nuclei develop along unique trajectories, some of which diverge from predictions of developmental theory. We find that those thalamic nuclei which show the most protracted development are at the greatest risk for later clinical differences in schizophrenia. The spatial pattern across thalamic nuclei for early psychosis risk is associated with a unique neuroreceptor fingerprint with implications for symptom severity.

## Main Text

The thalamus is a unique structure within the human nervous system, a major integrative hub home to one of the widest variety of synaptic inputs and projecting reciprocally to every region of cortex^1–3^. Given that a majority of its axonal connectivity originates from cortex^4^, and many cortical regions project to distal cortical regions via trans-thalamic connectivity^5^, the thalamus is situated as a key player in the global processing of nervous system information. Despite its central role in coordinating the vast array of sensory and cognitive processing, how its fine-scale tissue structures emerge from childhood into adulthood remain unknown within the human brain. The co-evolution of the thalamus with neocortex necessitated a relative expansion of higher-order cognitive representations in humans compared to animal models where much of the developmental neuroscience on the thalamus is derived. Coupled with the extremely protracted development of the brain in humans, and the suspected role of the thalamus in neurodevelopmental disorders such as schizophrenia^6–8^, the lack of a typical portrait of thalamic structural development beyond volumetric properties is a fundamental gap in human neuroscience. Recent volumetric studies find reduced volume in association nuclei of the thalamus in youth, which stabilizes in adulthood, and again returns to a negative trajectory in later life; sensorimotor nuclei are found to have stable volumes until later life^8^. Other studies have found increases in thalamic volume in childhood and declines in early adulthood^9–13^. In schizophrenia rather than early psychosis (EP), recent studies have found decreased volume of the mediodorsal, pulvinar, ventrolateral, ventral anterior, and ventral posterolateral thalamic nuclei and an association between reduced pulvinar volume and cognition^8,14–16^. Other studies have shown no difference in lateral nuclei volumes in early psychosis while differences exist in chronic schizophrenia; no association between nuclei volume and cognition in early psychosis; and no association between reduced volume and positive and negative symptoms in either early psychosis or chronic schizophrenia ^17^. Contradicting that, some research indicates pulvinar, mediodorsal, and ventrolateral volume deficits associated with early psychosis ^18^. Studies have provided conflicting evidence for specific involvement and direction of abnormality at the level of thalamic nuclei. In addition to cohort heterogeneity impacting the consistency of findings, volumetric measures do not index tissue properties with any sensitivity to the fine-scale structures we know to be developing. The question thus remains what is occurring at the level of tissue microstructure in development and disease in the thalamic nuclei?

Establishing a model for normative thalamic development, especially one that can be used to understand how its development has gone awry in neurodevelopmental disorders, requires large datasets with measures sensitive to the microstructural tissue properties that we know to be developing at the cellular level in the human thalamus^19,20^. Overcoming this hurdle, we combine here several large neuroimaging datasets with independent but complimentary measures sensitive to neurite and dendrite density^21,22^, as well as macromolecular tissue content^23^. Spanning the ages of 5 to 100+ years old, we chart the structural development of 44 thalamic nuclei (22 per hemisphere) ranging from sensory relay points to nodes of higher-order processing. In so doing, we will adjudicate between two hypotheses about the thalamus’ development. First, given the majority of axonal connectivity within the thalamus originates from cortex, thalamic nucleus development may inherit the well-described sensory-association axis of cortical development with sensory regions maturing before higher-order tissue. Alternatively, given the large convergence of connectivity from multiple cortical regions into a single thalamic nucleus^5^ and the unique cytoarchitecture of the thalamus^24,25^, the thalamus may diverge from the sensory-association axis observed in the cortex and cerebellum^26,27^. In the process, we demonstrate a largely anterior-posterior spatial tissue gradient in the thalamus validated across two quantitative measures. We apply our normative modeling to early psychosis to characterize the microstructural differences and their overlap with development, direction of EP developmental abnormality, and ML-assisted prediction of EP status from development. Finally, we investigate possible molecular mechanisms that may be associated with EP risk, demonstrating an overlapping spatial topography between receptor/transporter system organization and EP risk, and relating this alignment to symptom severity in early psychosis.

## Results

We first sought to characterize the spatial topography of tissue properties across the adult thalamus, given that inter-areal differences in neocortex form a sensory-association gradient in adults^28^. In n=1287 adult participants (ages 17 - 39) from the HCP datasets, we segmented the thalamus into 44 of its constituent nuclei^29^ using T1-weighted structural MRI (**Fig 1a**). In each nucleus, we can extract a structural metric sensitive to the density of neurite and dendrites^21,22^ by taking the ratio of T1 and T2-weighted images in each participant. Qualitatively, when rendered in their native volumetric space and colored according to neurite density, a clear gradient emerges across thalamic nuclei, oriented along an anterior-posterior axis (**Fig 1b**). While this data has the advantage of its large sample size, the T1/T2 ratio is a unitless measure. Can we replicate this effect with a quantitative metric that more directly measures fine-scale tissue properties? Recent advances in quantitative MRI^23,30^ have made possible the in vivo characterization of both the quantity and content of microstructural tissue properties. One such property, referred to as proton relaxivity (R1) quantifies the rate (1/sec) at which protons relax to field alignment, and is sensitive to both the quantity (milliliters) and composition of tissue within a voxel, scaling linearly with the presence of hydrophobic material such as myelin. Quantitative MRI was performed on N=79 participants ranging in age from 5 to 28 years old, whose thalamic nuclei were segmented through the same pipeline as HCP subjects. In qMRI participants 17 years of age and older, we extracted the mean R1 value from each nucleus in each participant, and when averaging across participants, we find a nearly identical gradient across the volume of the thalamus (**Fig 1c**). Correlating the T1/T2 ratio of each nucleus from the HCP data with the relaxivity measures from the qMRI data, we find that the two measures are significantly correlated with one another (**Fig 1d**; r = 0.92, p < 0.001).

**Figure 1:**
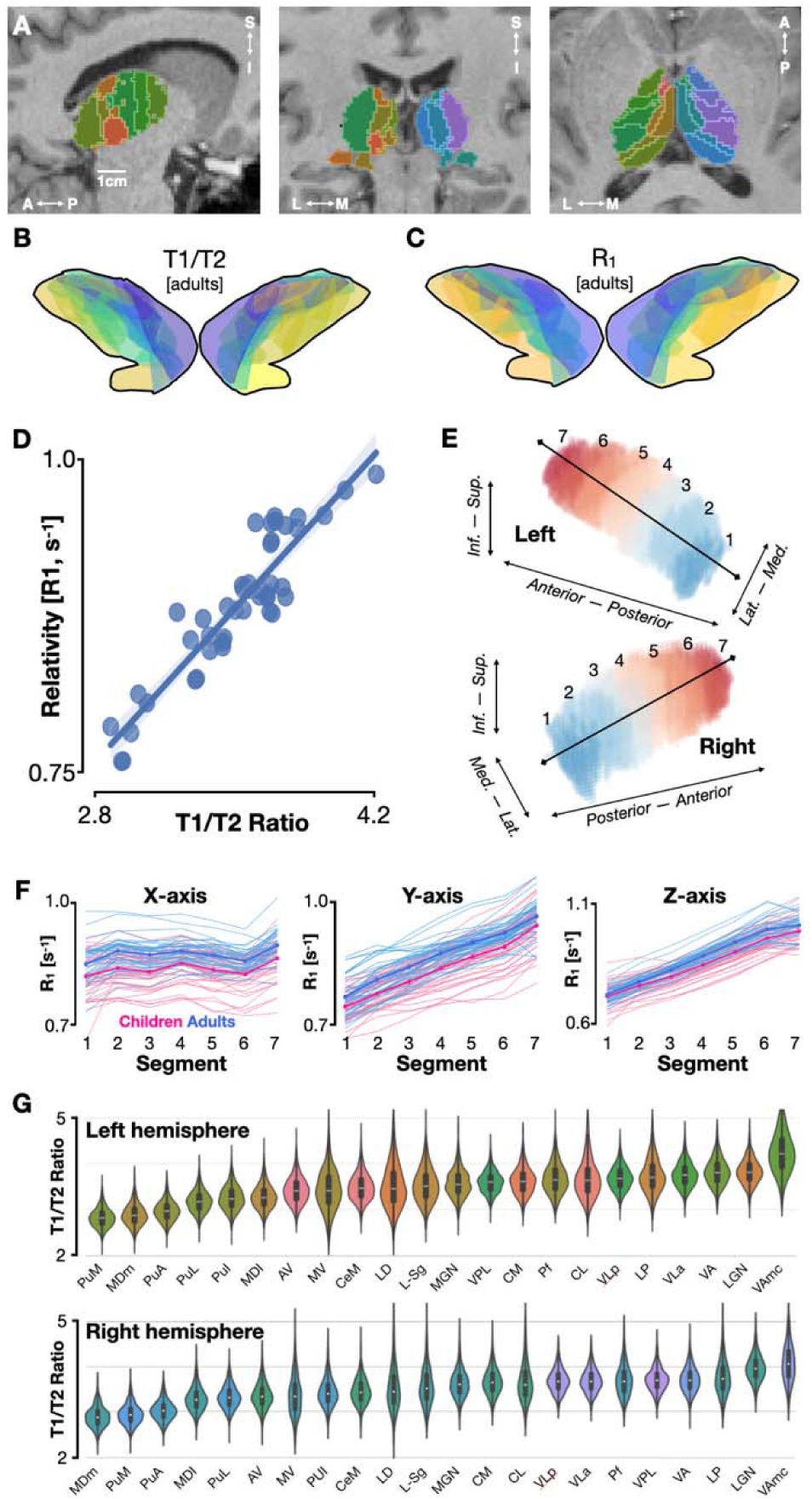
A replicable gradient of tissue microstructure across human thalamic nuclei in adults and children. **(A)** Segmentation of an example adult thalamus into its 22 constituent nuclei in both hemispheres. Blue tones color right hemisphere nuclei, green tones color left. **(B)** 3D rendering of the nuclei composing the adult human thalamus colored by mean T1/T2 ratio value. **(C)** Same as panel B but nuclei are colored by mean relaxivity across adults as measured by qMRI in a separate group of participants. A similar gradient is visible. **(D)** Scatterplot illustrating strong correspondence across thalamic nuclei between their mean T1/T2 ratio (x-axis) and their mean relaxivity (R1, y-axis). Each dot is a nucleus, values are averaged across voxels within a participant, then across participants. **(E)** A data-driven derivation of the principle axis (black line vector) along which relaxivity values vary most strongly across participants. Thalamus is segmented into seven equally spaced bins along this axis. **(F)** Values within each anatomical bin and each participant shown along the X, Y, and Z axes of the vector derived and illustrated in panel E. Children are shown in pink and adults are shown in blue. Both groups show similar and graded changes in relaxivity (R1) primarily along the Y and Z axes of neuroanatomical images. **(G)** Violin plots illustrating the mean T1/T2 ratio across participants, rank d from least tissue-dense on the left to the densest on the right. Nuclei are colored according to those shown in panel A.

To better quantify the shape of this thalamic gradient, and ensure that its presence is not an artifact of the way that the thalamus is segmented into nuclei, we use a data-driven approach which considers all voxels within an individual’s thalamus and tests for graded changes along many 3-dimensional vectors^31^(Methods). The axis describing the largest variance in qMRI tissue properties is oriented largely anterior-posterior with an upward trajectory as one travels anterior along the hemispheres of the thalamus (**Fig 1e**). This gradient is very similar in adult and child groups, who both show increasing tissue density as one travels from the ventral-posterior thalamus to the dorsal-anterior extent (**Fig 1f**). To quantify if the tissue environments across adult thalamic nuclei are statistically distinct, we performed a repeated-measures ANOVA as a function of thalamic nuclei in the T1/T2 ratio data of young adult participants. We find in both the left and right hemisphere that tissue density varies significantly between nuclei (left: F=2961, p<0.001, right: F=2431, p<0.001). Ranking thalamic nuclei from least to most tissue-dense leads to largely similar orderings between hemispheres, with sensorimotor nuclei such as the LGN, VA, and VPL ranking among the most tissue-dense (**Fig 1g**; Spearman rank coefficient ρ=0.16, p>0.05).

While previously observed in neocortex^32–34^ and recently in the cerebellum^26^, it remains to be empirically demonstrated whether the fine-scale gradient of tissue density within the thalamus reflects the intrinsic rate at which these regions develop during childhood. That is, is the sensorimotor axis of thalamic tissue density observed in adulthood related to each nucleus’ rate of development? To address this question, we repeated our thalamic parcellation in N=458 children aged 5-16, all of whom underwent structural MRI similar to adults to derive brain-wide maps of fine-scale tissue density as measured by T1/T2 ratio (Methods). When comparing tissue density between children and adults, we find that there is variability across thalamic nuclei in the magnitude of development. Interestingly, both low-level sensorimotor nuclei such as the LGN and higher-level association nuclei including inferior and lateral pulvinar show dramatic proliferation of tissue into adulthood (Fig 2a). When visualizing nuclei according to their development magnitude in R1 values, it is clear that the thalamus’ childhood development diverges from the tissue density gradient observed in R1 adult values (Fig 2b). When correlating adulthood R1 values and development magnitude, we observe no significant correlation (Fig 2c), suggesting thalamic tissue proliferation follows a unique routine compared to neocortex and the cerebellum. We observe that the patterns of thalamic development do not differ between hemispheres, with the association nuclei LP, LD, and pulvinar ranking among those with the greatest rate of development (Fig 2d; Spearman rank coefficient ρ=0.24, p>0.05). Developmental trajectories for specific nuclei that show small, medium, and large developmental effects show similar patterns when measured with qMRI (Fig 2e, see Fig S1-S2 for all nuclei). Thalamic nuclei proliferating tissue at unique rates would suggest that the broader-scale topology of fine-scale tissue properties across the thalamus is changing. Indeed, we find that development entails thalamic nuclei becoming more differentiated with age, with variance in tissue properties across nuclei significantly increasing with age (Fig 2f).

**Figure 2:**
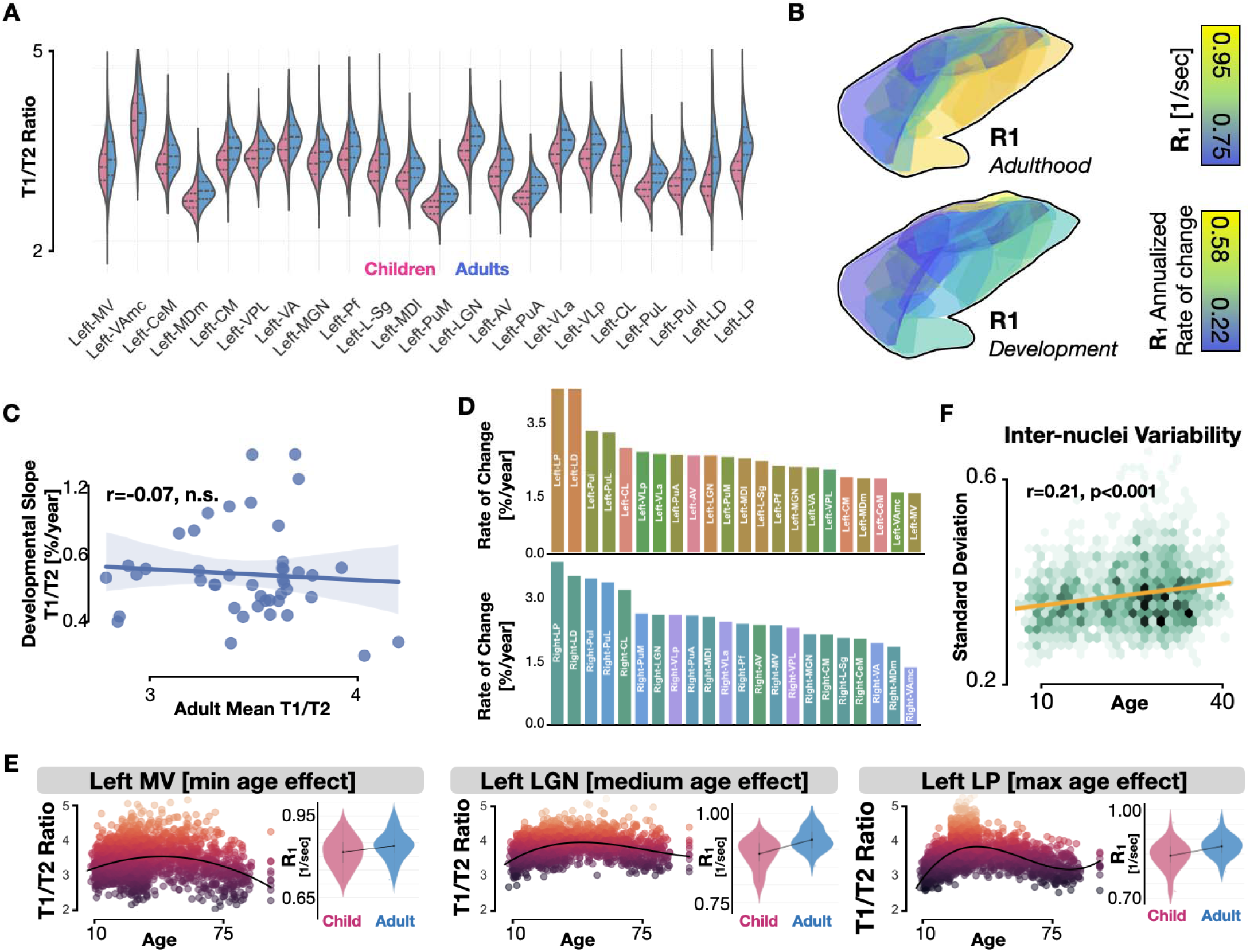
Protracted development of thalamic tissue microstructures diverges from adulthood tissue gradient. **(A)** Split violin plots comparing children and adult T1/T2 ratio values in each nucleus of the left thalamus. Mean T1/T2 ratio value is extracted per participant, violin depicts the distribution across participants. Nuclei ranked left to right in ascending effect size. **(B)** Nuclei of the thalamus (view of right hemisphere) colored by either adulthood relaxivity values (top) as measured by qMRI, or the group difference of relaxivity from childhood to adulthood. **(C)** Scatterplot comparing the adulthood mean T1/T2 ratio value of each nucleus with its annualized rate of development of T1/T2 ratio across childhood. Line of best bit and shaded error region depict bootstrapped linear regression which shows no significant correlation. **(D)** Annualized rate of change in T1/T2 ratio of each nucleus ranked from largest to smallest developmental effect. Top bars represent left hemisphere nuclei, bottom bars right hemisphere. **(E)** Example scatterplots of T1/T2 ratio as a function of age in three thalamic nuclei with low (MV), medium (LGN), and high (LP) developmental effect sizes. Each dot represents the mean T1/T2 ratio value in a given participant’s nucleus in the left hemisphere. Violin plots on the right depict the mean relaxivity (R1) in the same nucleus as measured by qMRI in an independent set of participants, violin width denotes participant density. **(F)** Binned scatterplot across all child and adult HCP datapoints depicting the standard deviation of T1/T2 ratio values across thalamic nuclei of both hemispheres plotted against each participant’s age. Orange line denotes line of best fit.

With a normative model of both the distribution and development of fine-scale tissue properties across the human thalamus, can we begin to characterize how development is atypical in neurodevelopmental disorders such as schizophrenia? While such a condition can be heterogeneous in its symptoms and treatment trajectory–conditions which can confound our ability to measure replicable biomarkers–brain-based measurements from individuals shortly after their first episode but before medication (early psychosis: EP) can provide a relatively unadulterated picture of the nervous system in schizophrenia. To that end, we parcellated the thalamus and aligned images of T1/T2 ratio from a large dataset of EP diagnosed patients (N=110) and matched neurotypical (NT) participants (N=58). Extracting the mean T1/T2 ratio from voxels within each thalamic nucleus, we find that differences between NT and EP participants vary by nucleus (**Fig 3a-b**). Nuclei including right AV show less tissue density in EP individuals, while others such as right PuL seem to show similar values (**Fig 3d**). A 2-way mixed model ANOVA with factors of diagnosis and nucleus revealed significant main effects for diagnosis (left: n.s.; right: F = 13.5, p < 0.001) and nucleus (left: F = 3225, p < 0.001; right: F = 2674, p < 0.001) as well as a significant interactions between diagnosis and nucleus (left: F = 18.4, p < 0.001; right: F = 21.3, p < 0.001). When visualizing the top 50% of nuclei that show the largest absolute difference in tissue density between EP and NT participants (**Fig 3a**), we find that these nuclei largely overlap with the top 50% most developmentally dynamic nuclei. Indeed, correlation of each nucleus’ rate of development with the magnitude of its tissue density difference between EP and NT groups yields a significant correlation (r = 0.52, p < 0.001; **Fig 3c**). Because fine-scale tissue properties appear to be less dense in EP individuals, the ontogeny of this difference remains an open question. Indeed, both younger children and aging adults both show less tissue density compared to young adults (**Fig 2e; Figs S1-S2**). Did tissue fail to proliferate during earlier development, in which case the thalamic environment of EP patients may better resemble children, or has a degenerative loss of fine-scale structures in EP led to a thalamic environment more consistent with older individuals?

**Figure 3:**
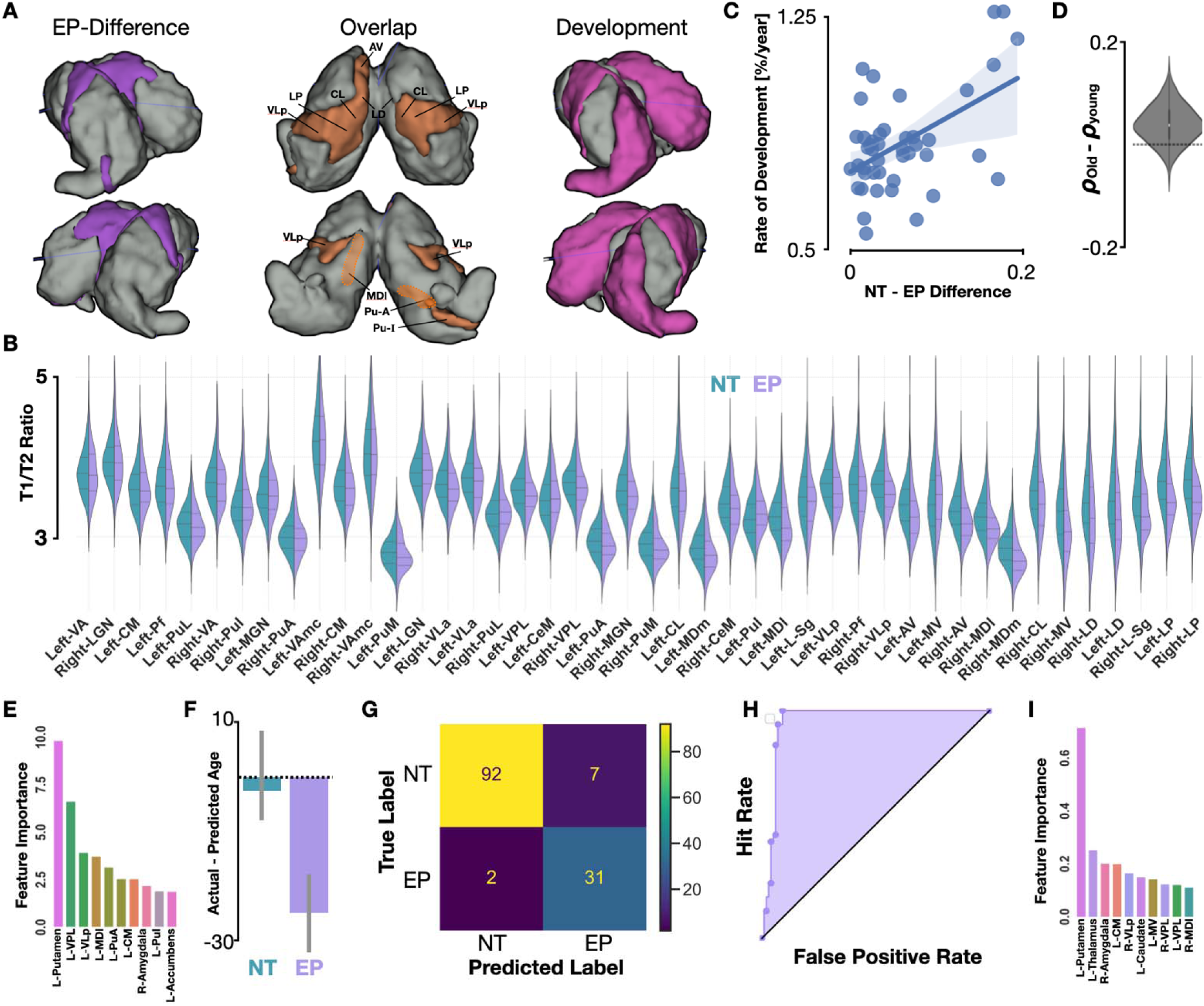
Thalamic microstructural tissue differences in early-psychosis schizophrenia. **(A)** Top 50% of nuclei for those with largest effect size for EP diagnosis compared to neurotypical (left, purple), top 50% of nuclei showing largest developmental effect across childhood (right, pink), and those nuclei which overlap between these two sets (middle, orange). **(B)** Split violin plots comparing the T1/T2 ratio across neurotypical (NT) and early-psychosis schizophrenia (EP) diagnosed individuals. **(C)** Significant correlation between the diagnostic difference of a given nucleus’ T1/T2 ratio difference between neurotypical and EP diagnosed adults (x-axis) and the rate of that nucleus’ development during childhood in neurotypical children. **(D)** Violin plot depicting the distribution across EP patients when the T1/T2 ratio of thalamic nuclei is correlated with the average T1/T2 ratio of nuclei from children or aging adults of the HCP datasets, taking the difference between resulting Pearson coefficients (y-axis). Distribution is significantly above zero, suggesting EP thalamic nuclei tissue topography resembles older participants. **(E)** The importance of the top anatomical features in the prediction of age extracted from the Lasso regression model. **(F)** A Lasso regression model results incorporating polynomial features trained to predict an individual’s age through a 10-fold CV approach shows no significant deviation from actual age in neurotypical individuals but produces significantly older predicted ages for adults diagnosed with early psychosis. **(G)** Confusion matrix for a linear support-vector classifier trained to classify NT from EP patients shows high accuracy and recall performance. In addition to thalamic nuclei, other subcortical structures are included in this model such as putamen, amygdala, caudate, and accumbent. **(H)** Receiver-operating characteristic (ROC) curve illustrating the hit and false-alarm rates of the support vector classifier. **(I)** The importance of the top anatomical features to classification performance from the diagnosis Lasso regression model.

To that end, we trained a Lasso regression model for brain-age prediction which uses mean T1/T2 ratio of each of the 44 nuclei across the left and right hemispheres as well as other subcortical structures, specifically each of the left and right caudate, putamen, accumbens, pallidum, amygdala, and gross thalamus as 12 additional inputs to define a relationship between subcortical nuclei and a participant’s age. Development of these additional subcortical structures is illustrated in **Figs S1-S2**. These other subcortical nuclei were included given heavy connectivity with thalamic structures^25^. Importantly, we include neurotypical individuals ranging in age from 5 to 100 in order to more widely estimate the relationship between the subcortical tissue environment and age. This model was capable of explaining 72% of the variability in age, with weights spanning thalamic, striatal, and amygdalar structures (**Fig 3e**). We find that EP patients have a predicted age significantly older than their true age based on subcortical tissue properties, while NT participants yield predicted ages consistent with their true age (**Fig 3f**). Extraction of the top 10 feature weights contributing to model performance revealed that the left putamen, VPL, VLp, MDI, and PuI were highly predictive of age (**Fig 3e**). To verify this model, we produced two average maps of T1/T2 ratio in the thalamus of NT participants: one from participants 10 ± 2 years younger than the youngest EP patient, and one from participants 10 ± 2 years older than the oldest EP patient. In each EP patient, we derived the mean T1/T2 ratio of each thalamic nucleus and performed a Pearson correlation with the mean values derived from each of the NT maps. For each EP patient, we derived the difference between the two resulting Pearson’s r values (old minus young), and we found that EP participants show a significant difference (**Fig 3c**; t = -8.42; p < 0.001), with their pattern of tissue density across thalamic nuclei better matching that of older participants. Given these differences, a linear support vector classifier can significantly classify EP from NT participants using T1/T2 properties of subcortical nuclei with an F1 score of 87% and accuracy of 93% (**Fig 3g**). A receiver operating characteristic (ROC) curve revealed an area under the curve (AUC) of 0.95 (**Fig 3h**). Features with the highest weights (**Fig 3i**) include left putamen, left whole thalamus, right amygdala, and thalamic nuclei such as left CM, right VLp, and left/right VPL.

We next sought to ground our development and early psychosis findings in receptor/transporter systems in order to identify possible molecular mechanisms of typical development and EP risk. We used PET-derived neurotransmitter system maps of receptors/transporters (Methods) to extract a proxy of the mean density of each of 22 molecular targets within each thalamic nucleus. We found that the developmental rate of a nucleus correlated with the density of 10 separate molecular targets. The strongest associations that survived FDR correction (p<0.05) when using a BrainSMASH spatial null were observed for 5HT1b (r=-0.724), H3 (r=-0.694), D2 (r=-0.585), FDOPA (r=-0.583), NET (r=-0.583), 5HT1a (r=-0.538), 5HTT (r=-0.535), NMDA (r=0.508), 5HT4 (r=-0.392), and MU (r=-0.367), which show a spatially aligned profile of neurotransmitter systems with that of structural developmental as assessed with MRI (**Fig 4a**).

**Figure 4:**
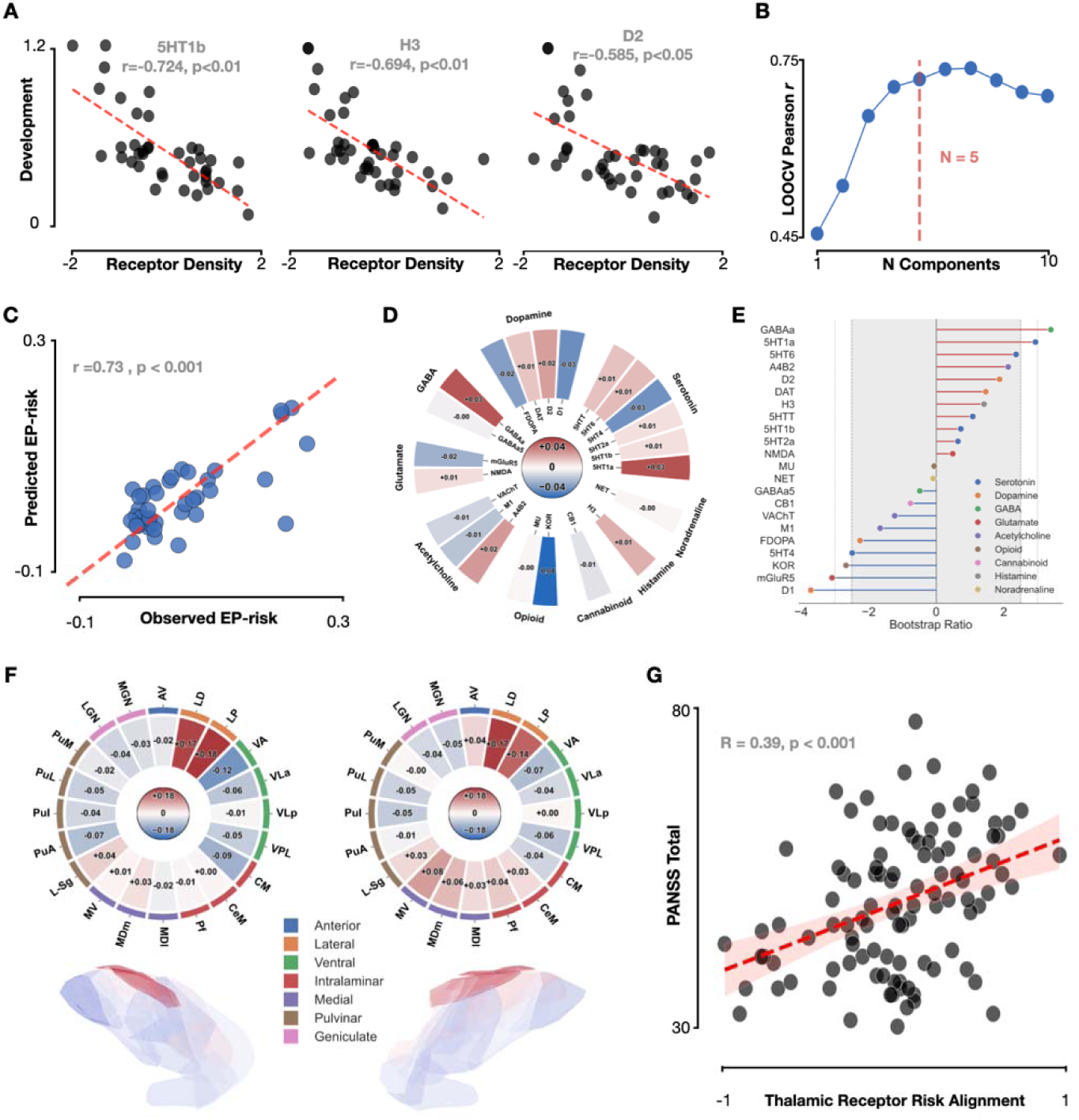
The Receptor/Transporter System Fingerprint of Thalamic Early Psychosis Risk. **(A)** Top three correlations between receptor density and development rate of T1/T2 values across thalamic nuclei. **(B)** Partial least squares regression model performance as a function of component number, assessed using a leave-one-out cross validation procedure. Model performance plateaued at N=5 components. **(C)** PLS model predictions for a given thalamic nucleus’ structural EP risk scattered against the observed structural EP risk for that nucleus. Results show a significant correlation, Pearson coefficient and p-value shown with inset text. **(D)** The receptor weights for each receptor type from the PLS regression model grouped by receptor class (outer ring text). Colorbar at center denotes weight value. **(E)** Lollipop plot depicting the reliability metric for each receptor evaluated using a bootstrapped approach in the PLS regression model. **(F)** Nucleus scores for each nucleus shown in polar heatmaps for the left and right thalamic nuclei. Nucleus families shown in coloring of outer ring, and inset text shows family labels. Lower renderings color each thalamic nucleus using the same color scale in the polar heatmaps. **(G)** The more an individual’s T1/T2 structural pattern across nuclei aligns with the PLS-derived pattern, the worse their symptom severity as measured by PANSS when controlling for antipsychotic exposure.

Interestingly, despite our findings of EP pathology correlating with rate of development, we identified no single molecular target associated with the spatial profile of structural EP-NT difference across nuclei. We further found that development and EP pathology only minimally overlapped in their molecular associations (Spearman coefficient = 0.102, p > 0.05). The nuclei that are most at risk in EP are heterogeneous, and EP pathology may impact nuclei which have unique receptor profiles rather than nuclei aligned with any single receptor system. We thus used PLS to derive latent variables from the thalamic neurotransmitter system maps most predictive of NT-EP difference, resulting in a 5-component model with LOOCV out-of-sample performance significant under a conventional permutation test (r = 0.73, p < 0.001) and still significant under a spatial null test that addressed autocorrelation (r = 0.73, spatial null p < 0.05; **Fig 4b-c**). The performance of this model was largely preserved with a stricter nested spatially blocked LOOCV procedure (0.56). We found that the model predicted the highest nucleus scores for the left and right LD and LP nuclei (0.14-0.18). Nuclei with higher receptor-derived PLS scores have receptor/transporter profiles that are more predictive of greater EP risk (**Fig 4f**). The receptor/transporter coefficients of this model indicated that the strongest contributors by absolute coefficient magnitude were KOR, GABAa, and 5HT1a (**Fig 4d**). Bootstrap ratios were computed, revealing 5 receptor targets that are reliable contributors anchoring this latent neurochemical profile: D1 (BSR = −3.71), GABAa (+3.39), mGluR5 (−3.08), 5HT1a (+2.94), and KOR (−2.67) (|BSR|>=2.5; **Fig 4e** and Supplementary Table 2 for the full list ^35^).

To investigate the subject-level alignment to the receptor/transporter-derived PLS profile, we calculated for each subject an alignment score which expresses how strongly the spatial pattern of T1/T2 ratio values across nuclei aligns with the receptor/transporter-derived PLS profile by projecting each subject’s thalamic tissue profile onto the PLS profile. A subject with higher score would have a tissue pattern that more strongly matches the receptor-derived PLS profile that is associated with the NT-EP tissue difference map. We found that the extent to which a subject’s thalamic tissue pattern resembles the receptor-driven pattern that characterizes early psychosis risk is associated with psychotic symptoms. Specifically, receptor score, controlling for antipsychotic (CPZ) exposure, predicts psychotic symptoms, specifically the Positive and Negative Syndrome Scale (PANSS; clinician administered) total score (**Fig 4g**; p < 0.001) as well as positive (p < 0.01) and general (p<0.001) psychotic symptom subscales. This relationship does not extend to negative psychotic symptoms as measured by PANSS or Clinical Assessment Interview for Negative Symptoms (CAINS) or to depression or mania symptoms. To confirm that the relationship was attributable to receptor weighting rather than to overall tissue, the mean of the z-scored tissue across nuclei was used as the independent variable, which showed that these findings were not driven by absolute tissue quantities (p > 0.05) but rather the spatial profile. Thus, these data suggest that individual variation in the extent to which the pattern of fine-scale thalamic tissue aligns with the PLS-derived receptor/transporter pattern is associated with with positive and general psychotic symptom severity in early psychosis.

## Discussion

We find that thalamic nuclei show distinct fine-scale tissue environments measurable in the living brain using both standard and quantitative MRI techniques. The thalamus demonstrates a spatial gradient of relaxivity in the adult brain (**Fig 1**), suggesting an axis of decreasing tissue density oriented largely anterior-posterior. Surprisingly, this spatial gradient of tissue density does not correspond with a spatial gradient of tissue development during childhood (**Fig 2**), suggesting the thalamus follows a developmental routine distinct from cortex and the cerebellum. Highly myelinated sensory nuclei, such as the LGN, demonstrate a proliferation of fine-scale tissue structures as large and protracted as that of those considered association nuclei such as the pulvinar, even when quantifying development by normalizing for initial tissue density (**Fig 2d**). Lastly, we find that thalamic nuclei show significantly less fine-scale tissue in those diagnosed with early psychosis schizophrenia compared to neurotypical individuals (**Fig 3**). Those subcortical nuclei which show the largest difference between NT and EP individuals tend to be those which show the most dramatic proliferation of tissue during development. Overall, these results provide a normative portrait for the distribution of fine-scale tissue properties across the thalamus and how they develop from childhood into adulthood, providing the groundwork for understanding how development is atypical in neurodevelopmental disorders such as schizophrenia.

While in neocortex and the cerebellum^26,27,36^, there appears to be a relationship between a brain region’s tissue density (as measured with T1/T2 ratio) and its developmental rate, this relationship does not appear to be a universal principle of the central nervous system, with thalamic nuclei deviating from this pattern. The LGN, for example, a region heavily interconnected with primary visual cortex, is largely considered a nucleus dedicated to the relay and feedback of primary sensory information, but shows dramatic growth of fine-scale tissue structures from childhood into adulthood where prior models would have predicted little growth. Thalamic nuclei also become more structurally distinct from one another with development (Fig 2f), which differs from the cerebellum whose constituent lobules become more similar across development^26,27,36^. One possible explanation for the unique patterns of development in thalamic nuclei may result from the fact that a given nucleus can send projections which span multiple low- and higher-level cortical regions as observed in humans with functional MRI^37^. Furthermore, it has been observed that connectivity patterns of cortex with subcortical regions can change sign (excitatory versus inhibitory) with development^38^, suggesting there may not be a simple linear mapping between cortical development and subcortical structures such as the thalamus.

Our results relate to prior observations of a connectivity gradient in the human thalamus^39,40^, but we extend these findings to demonstrate a fine-scale tissue gradient across nuclei. In regard to the atypical development of thalamic tissue content in EP, we find that while fine-scale tissue structures appear to be less dense in some thalamic nuclei of early psychosis patients, the overall distribution of tissue content across thalamic nuclei in EP patients appears more consistent with older brains compared to younger brains, suggesting that the neuroanatomical origin of EP is not simply delayed or stunted development. The broad sparsity of fine-scale tissue is consistent with global cognitive impairments reported in patients with schizophrenia^41^. Our work also replicates and extends prior work finding volumetric reduction in higher-order thalamic nuclei in schizophrenia^18^, suggesting reduced fine-scale density of neurite structures may drive these gross volumetric differences as well as the thalamus’ impoverished functional connectivity to cortex^42^. Our ability to classify EP from NT participants with relatively high precision from fine-scale tissue properties of subcortical structures suggests future work should focus on the elucidation of these properties in subcortex through mapping the longitudinal development and function of schizophrenia.

While the use of independent normative PET datasets to derive molecular maps for spatial analysis of our findings is characterized by numerous limitations discussed below, we sought to ground our findings in a possible molecular mechanism of the spatial organization of the observed EP risk that we have presented. We found that there is a spatial topography of receptor/transporter density associated EP risk structural topology in early psychosis, linking molecular patterns to patterns of observed tissue abnormalities. Further, we found that PANSS positive and general subscale symptom severity could be predicted by a risk-predictive tissue pattern derived from a receptor/transporter density multivariate profile anchored by the D1, mGluR5, KO, 5HT1a, and GABAa receptors. These specific receptors have been broadly implicated in schizophrenia pathology; for example, D1 receptor density, which has been observed to be reduced in early psychosis in the prefrontal cortex, has been implicated in general and social cognition and positive symptoms ^43–45^. Availability of the KO receptor has been shown to predict anhedonia, selective antagonism of KOR to produce antidepressant effects, and selective agonism to promote depressive symptoms, which aligns with symptoms indexed by the PANSS general subscale (G6) ^46,47^. Our findings highlight the need for further research using quantitative MRI measures to investigate development in typical and EP populations and to directly link that development to molecular mechanisms, which is not possible for us to do in this dataset. Understanding the specific molecular mechanism associated with the observed tissue abnormalities across development may shed light on the etiology and neurodevelopmental trajectory of schizophrenia, providing insight into approaches in personalized medicine to improve treatment and diagnosis of schizophrenia spectrum disorders.

## Limitations and Future Directions

Replications and extensions of the current findings using larger quantitative MRI datasets are encouraged to further clarify the relationship between these two measures and their ability to shed light on development and pathology. The question of what T1/T2 ratio measures continues to be queried. Previous research has suggested that the MRI-based measure of T1/T2 ratio values may correspond to cortical myelin content as mapped with histology by Hopf^28,48^. Other research based on post-mortem histology and imaging^22,49,50^ suggests it may not track myelin, but instead that this measure corresponds to the density of neurites, and more specifically in some references dendrites, as well as reflecting iron density. Quantitative MRI, on the other hand, while not existing in large repositories like T1/T2 data, provides a more quantitative definition of the microstructural properties of the living brain while reducing the impact of scanner biases and iron content^23^. The strong correlation between qMRI and T1/T2 metrics in the current study offer converging evidence for protracted tissue development in the human thalamus and an atypical tissue sparsity in thalamic nuclei of those diagnosed with schizophrenia. Additionally, our findings regarding the thalamic environment of NT and EP individuals remain cross-sectional observations MRI data within multiple large datasets collected across different imaging collection and recruitment landscapes. Thus, future research should focus on investigation of microstructural development using quantitative MRI methods in longitudinal representative samples of individuals to observe patterns of development that precede eventual conversion to schizophrenia spectrum disorders. In addition, larger samples than those presented here may allow for finer discrimination between subtypes of psychotic disorders and their developmental neuroanatomical etiology.

While we investigated possible molecular mechanisms that may be associated with EP risk, our measures only permit analysis of overlapping spatial organization between receptor density and EP-associated tissue differences. We are further limited since the receptor/transporter maps used are sourced from several independent PET datasets rather than being from the same subjects on from whom our thalamic tissue values were extracted, precluding our ability to link individual subject-level molecular variables to behavior in any causal way. The question of how receptor/transporter systems may relate to individual variation in thalamic tissue microstructure in schizophrenia and early psychosis in development must be pursued in future research. Future longitudinal studies should track both quantitative tissue trajectories in early psychosis and controls as well as molecular targets in order to be able to provide direct evidence supporting a developmental relationship between EP pathology and molecular substrates.

## Methods

### Datasets and Subjects

**The Lifespan Human Connectome Project Development (HCP-D) 2021 release** contained data from 652 healthy subjects ages 5-21, of which 628 possessed all relevant data and were included in the analysis (mean=14.4±4.03; males=283). Briefly, as detailed in the previous literature^51^, the study recruited healthy participants who were proficient in English and could commit to completion of the study, with the aim of encompassing a broad cross-section of typical development. Exclusion criteria included learning disabilities, disorders of development, any health condition that is known to affect developmental trajectory, rare health conditions that would invalidate anonymization, and any contraindications of MRI. Structural MRI data were collected on a 3T Siemens Prisma platform. The Siemens 32-channel Prisma head coil was used for subjects ages 8-21 at the time of the scan and the Ceresensa pediatric 32-channel head coil was used for subjects ages 5-7. The high resolution T1- and T2-weighted scans were collected for all the subjects in this release following the protocol documented in previous publications^52,53^. Briefly, for the T1w protocol 0.8mm isotropic scans were collected using the 3D MPRAGE sequence (sagittal field of view (FOV)=256 x 240 x 166 mm; matrix=320 x 300 x 208; TR=2500 ms; TE=1.8/3.6/5.4/7.2 ms; TI=1000 ms; FA=8°, bandwidth (BW)=744 Hz per pixel) For the T2w protocol, scans were collected with 0.8mm isotropic resolution ((sagittal FOV=256 x 240 x 166 mm; matrix=320 x 300 x 208 sagittal slices in a single slab; TR=3200 ms; TE=564 ms; turbo factor=314; TI=1000 ms; FA=8°, bandwidth (BW)=744 Hz per pixel). The data were minimally processed according to the procedure in Glasser et al. (2013); briefly, the pipeline involved correction of MR gradient-nonlinearity-induced distortion, alignment of the T1w and T2w scans, B1 bias field correction, and registration of native space structural volumes to MNI space, FreeSurfer automated segmentation of the volume, reconstruction of the white and pial cortical surfaces, and folding-based surface registration to the FreeSurfer standard surface atlas.

**The Human Connectome Project Young Adult (HCP-YA) S1200 release** contained data from 1200 healthy subjects ages 22-35, of which 997 have all relevant imaging data (mean=28.7±3.69). Briefly, as detailed in the previous literature^54^, the study recruited healthy participants from a population of twins and non-twin siblings born in Missouri who were proficient English and could commit to completion of the study. Exclusion criteria included documented learning disabilities, disorders of development, neuropsychiatric and neurological disorders, chronic health conditions like diabetes and hypertension, and pre-term birth (34 weeks and 37 weeks for twins and non-twins, respectively). Structural MRI data were collected on a customized Siemens 3T “Connectome Skyra” at the study site at Washington University, using the standard Siemens 32-channel Prisma head coil. As previously detailed, two sets of high resolution T1- and T2-weighted scans were collected following the HCP protocol. Briefly, for the T1w protocol two separate 0.7mm isotropic scans were collected using the 3D MPRAGE sequence (field of view (FOV)=224 mm; matrix=320, 256; repetition time (TR)=2400 ms; echo time (TE)=2.14 ms; inversion time (TI)=1000 ms; flip angle (FA)=8°, bandwidth (BW)=210 Hz per pixel; echo spacing (ES)=7.6 ms). For the T2w protocol, two separate scans were collected using the variable flip angle turbo spin-echo sequence (Siemens SPACE) with 0.7mm isotropic resolution (FOV=224 mm; matrix=320, 256; TR=3200 ms; TE=565 ms; BW=744 Hz per pixel). The scan of highest quality was used. The data were minimally processed according to the procedure previously noted.

**The Lifespan Human Connectome Project Aging (HCP-Aging) 2021 release** contained data from 725 healthy subjects ages 36-100+, of which 678 have all relevant imaging data (mean=60±15.5; males=296). Briefly, as detailed in previous publications^55^, the study recruited healthy participants who were proficient in English and could commit to completion of the study across 4 sites with matched protocols. Exclusion criteria included any major diagnosed disease. Structural MRI data were collected on a 3 T Siemens Prisma platform, using the Siemens 32-channel Prisma head coil. The high resolution T1- and T2-weighted scans were collected for all the subjects in this release following the protocol documented in previous publications^52,53^. Briefly, for the T1w protocol 0.7mm isotropic scans were collected using the multi-echo MPRAGE sequence (sagittal FOV=256 x 240 x 166 mm; matrix=320 x 300 x 208; TR=2500 ms; TE=1.8/3.6/5.4/7.2 ms; TI=1000 ms; FA=8°, bandwidth (BW)=744 Hz per pixel). For the T2w protocol scans were collected with 0.8mm isotropic resolution (sagittal FOV=256 x 240 x 166 mm; matrix=320 x 300 x 208 sagittal slices in a single slab; TR=3200 ms; TE=564 ms; turbo factor=314; TI=1000 ms; FA=8°, bandwidth (BW)=744 Hz per pixel). The data were minimally processed according to the procedure previously noted.

The Human Connectome Project Early Psychosis (HCP-EP) 1.1 release contained data from 183 subjects ages 16-35; the present study used a subset of 169 subjects with minimally preprocessed structural MRI data available, of which 166 have all relevant data and were included in the present study (EP: mean=22.8±3.59, males=43; NT: mean=24.9±4.08, males=19). Briefly, as detailed in the HCP-EP 1.1 Data Release Reference Manual and the previous literature^56,57^, the study recruited patients with affective and non-affective psychosis with onset not more than 5 years previous to study enrollment as well as healthy control participants who were proficient in English across 4 early psychosis programs with matched protocols. The non-affective psychosis group (N=83) have a DSM-5 Schizophrenia Spectrum and Other Psychotic Disorders diagnosis, a category that includes schizophrenia, schizophreniform, brief psychotic disorder, schizoaffective, delusional disorder, and psychosis not otherwise specified. The affective psychosis group (N=28) have a DSM-5 diagnosis of a mood disorder with psychosis, which includes bipolar disorder with psychosis (including most recent episodes of depressed and manic types) and major depression with psychosis (single or recurrent episodes). The healthy control group (N=58) were screened out if they reported current use of psychiatric medications, first-degree relatives with a diagnosed schizophrenia spectrum disorder, bipolar disorder, current anxiety disorder, past anxiety disorder with duration and recency of 12 months or requiring medication, any of the psychiatric disorders for which the clinical groups were recruited, or a lifetime history of psychiatric hospitalization. Structural MRI data were collected on 3 T Siemens MAGNETOM Prisma scanners, using the Siemens Prisma 32-channel head coil or the 64-channel head and neck coil, with the neck channels turned off. The high resolution T1- and T2-weighted scans were collected for all the subjects in this release following the protocol documented in the literature^56^. Briefly, for the T1w protocol 0.8mm isotropic scans were collected using the MPRAGE sequence (sagittal FOV=256 x 240 x 166 mm; matrix=320 x 300 x 208; TR=2400 ms; TE=2.22 ms; TI=1000 ms; FA=8°, bandwidth (BW)=220 Hz per pixel). For the T2w protocol scans were collected using the variable flip angle turbo spin-echo sequence (Siemens SPACE) with 0.8mm isotropic resolution (sagittal FOV=256 x 240 x 166 mm; matrix=320 x 300 x 208; TR=3200 ms; TE=563 ms; turbo factor=314, bandwidth (BW)=744 Hz per pixel). The data were minimally processed according to the procedure previously noted.

Quantitative MRI Dataset. The qMRI dataset used in the present study was previously collected at Stanford University and published in previous studies^30,32^. Participants were screened to have normal or corrected-to-normal vision and no psychiatric history, and informed consent was obtained at the original study site. The sample includes 79 subjects grouped into children ages 5-12 (N=41; mean=8.10±2.27) and adults ages 17-28 (N=38; mean=23.6±1.81). The imaging protocol used was sourced from Mezer et al. (2013). Structural MRI data were collected on a 3T GE Discovery MR750 scanner (GE Medical Systems), using a phase-array 32-channel head coil. Four spoiled gradient echo (spoiled-GE) scans were collected at 0.8x0.8x1.0 mm resolution (TR=14 ms; TE=2.4 ms; FA=4°, 10°, 20°, 30°). Four spin echo inversion recovery (SEIR) scans were collected with echo planar imaging read-out (TR=3000 ms; TE=minimum full with 2x acceleration; TI=50, 400, 1200, 2400 ms; in-plane resolution=2x2 mm; slice thickness=4 mm). The data were processed using the mrQ v.2.1 (https://github.com/mezera/mrQ) processing pipeline in MATLAB, the methods of which has been detailed in prior literature^58^). Relaxivity (R1), or the rate of return of a water proton to realignment with the field as measured in Hertz, is calculated using the spoiled-GE scans with the SEIR scans to control for field inhomogeneities. R1 represents the proportion of water versus myelin present in a given voxel, as validated in the previous literature^23^.

### Data Processing

Whole brain T1/T2 ratio maps were computed as the ratio of the T1w and T2w images, representing a relative measure of myelin content sensitive to dendritic density^22^. The thalamus was segmented into 25 nuclei per hemisphere using a histologically-based probabilistic atlas implemented in FreeSurfer 7.0^29^. The reticular, paratenial, and paracentral nuclei were excluded from the analysis due to insufficient resolution to reliably identify voxels assigned to these regions in all participants. The remaining 22 nuclei per hemisphere were included in the analyses. Outliers were removed from each dataset when they were 3.5 or more standard deviations from the mean for that dataset. Since the data were collected from MRI scanners at several different sites, neuroCombat was used for multi-site harmonization to reduce the effects of scanner variability on measurement, which produce increased power and reproducibility^59^. Parametric adjustments were performed for mean adjustments without scaling. Empirical Bayes was not performed. The biological covariates to preserve across sites were sex, age, and diagnosis.

### Statistical Methods

Statistical analyses were completed using the Python package Pingouin^60^. Pearson’s correlation coefficient was used to quantify the relationships between continuous variables, such as when measuring the association between two tissue measures and between development rate and group mean differences. A repeated measures ANOVA was performed to test for differences across nuclei tissue measures in young adults. Spearman’s rank correlation coefficient (rho) was used to assess the hemispheric laterality of the thalamic tissue patterns. A 2-way mixed model ANOVA with factors of diagnosis and nucleus was performed to examine main effects and their interaction.

To calculate the annualized percent change (APC) for each nucleus the percent change from the starting mean from ages 5-11 to the ending mean from ages 17-23 was divided by the year span separating the midpoints of each of those ranges. This scale-independent approach was used to account for different baseline values across variables. The APC formula is as follows:

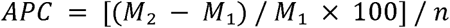

where M₁ = starting mean, M₂ = ending mean, and n = number of years.

### Developmental Trajectories in Typical Development

To generate developmental trajectories for thalamic nuclei, leave-one-out cross-validation (LOOCV) was used to fit polynomial models up to the 4th degree to each nucleus and select the polynomial model that minimized the mean square error as the best model. The selected polynomial model for each nucleus was then fit to the entire dataset. Visualizations of trajectories across the lifespan were generated with these parameters for each nucleus.

### Developmental Gradients in qMRI

The principle axis along which relaxivity values vary most strongly within the thalamus was derived using the mrGrad toolbox in MATLAB; for in-depth description, see the GitHub repository (https://github.com/MezerLab/mrGrad) and previous literature introducing the tool^31^. Briefly, this data-driven approach calculated the single vector decomposition of the coordinates specified in the default FreeSurfer volumetric masks of the left and right thalami, thereby delineating the principal orthogonal axes of the provided region. The thalamic volume mask is then divided into n equally-spaced segments by defining hyperplanes at either extreme of the data and placing n-1 parallel hyperplanes spaced at equal intervals between them and assigning each voxel in the mask to a segment based on distance. The median relaxivity value in each segment was calculated across the entire qMRI sample in order to reveal the potential gradients along the three principal axes. Further, we averaged these median values across subjects within the child and adult groups in order to allow comparison of the spatial gradients. To visualize group differences, the group-averaged data were plotted for each of the main orthogonal axes X, Y, and Z.

### Diagnosis Prediction

The scikit-learn Python module was used to perform the processing and analysis for machine learning classification in this study^61^. The aim of this analysis was to perform binary classification of HCP subjects into early psychosis versus healthy control participants using the T1/T2 ratio values previously extracted from the subcortex. All HCP datasets except for the HCP-EP dataset were collated into one dataset. Due to the massively imbalanced nature of the dataset as well as the expanded age range of 5-100 for the control dataset compared to 16-35 for the EP dataset, we performed binned random under-sampling from this combined control dataset without replacement in order to create a new sample of control participants that conformed to the same distribution of ages as the HCP-EP dataset in order to provide better correspondence between healthy controls and EP patients and reduce bias^62^. The HCP-EP data were combined with the undersampled data (EP=110; NT=329), resulting in a significantly less imbalanced dataset at a ratio of 1:3 compared with 1:12 in the overall dataset. This procedure was conducted k=10 times and the average mean performance of each model as detailed below across each subsample was calculated for use in model selection.

The dataset was split into features and labels. The features comprised the 44 bilateral thalamic nuclei as well as the bilateral whole thalamus, caudate, putamen, pallidum, accumbens, and amygdala, totaling 56 features in all. The labels were the binary diagnostic categories of control subjects and early psychosis subjects, the latter of which combined both the non-affective and affective psychosis groups since there were a more limited number of subjects in each of these subgroups. Using the train_test_split function, the test and train datasets were shuffled and divided in a 70/30 split with stratification to ensure class distribution in the full dataset is reflected in the test and train datasets, considering this imbalanced dataset includes fewer early psychosis subjects than healthy controls. Standardization of the features of the dataset to zero mean and unit variance was performed using StandardScaler. The features of the train dataset were used to fit the StandardScaler object, which was then used to standardize both the train and test data.

For model selection, the following models were evaluated: logistic regression, random forest, support vector machines (SVM), and linear SVM. Briefly, each model was initialized into a pipeline object; parameter grids were defined for each to optimize based on that set of hyperparameters; tuning of the hyperparameters was performed with GridSearchCV using the F1 score to assess performance since this imbalanced dataset would benefit from a scoring metric that balances precision and recall by not weighting the score for correct detection of the majority negative class (no early psychosis diagnosis) while prioritizing correct detection of the minority positive class (early psychosis diagnosis); and cross-validation of this process was performed with cross_val_score using a stratified K-fold procedure in order to ensure class distribution for each training fold does not significantly deviate from the overall dataset, resulting in k=5 folds for the inner grid search cross-validation, then k=10 folds for the outer cross-validation, with the mean F1 score of those five k=5 grid search cycles used to determine the best performing model.

This procedure identified the linear SVM model as the best performing model (hyperparameters: C=0.1, tolerance=0.1, penalty: L1). The mean performance of the model on training data during model selection as measured with the F1-score metric was 84%. Finally, the best model selected through the above process was retrained on the full train dataset (F1 score=88%). The test split was used to assess the final classification performance on out of sample data with this model. In order to visualize and better interpret the performance of the selected model in discriminating between the two classes, a confusion matrix was generated. A receiver-operating characteristic (ROC) curve was plotted to visualize the true positive versus false positive rate of the selected model compared to a no-skill line.

### Age Regression

The aim of this analysis was to create an age prediction model trained on a control dataset to compare the age predictions of the model for an out of sample control dataset and an EP dataset. The procedure for this process was similar to that used previously for the classification of these data. Briefly, the T1/T2 ratio values previously extracted from the subcortex were used as the features. All HCP datasets (developmental, young adult, and aging) except for the HCP-EP dataset were collated into one dataset. The entirety of the dataset was retained in order to maintain information for prediction of ages across the lifespan. The dataset was split into features and labels. The features comprised the 44 bilateral thalamic nuclei as well as the bilateral whole thalamus, caudate, putamen, pallidum, accumbens, and amygdala, totaling 56 features in all. The labels were the ages of the subjects. Using the train_test_split function, the test and train datasets were shuffled and divided in a 70/30 split. Standardization of the features of the dataset to zero mean and unit variance was performed using StandardScaler. The features of the train dataset were used to fit the StandardScaler object, which was then used to standardize both the train and test data.

For model selection, the following models were evaluated with and without polynomial (2nd degree) features: linear regression and lasso regression. Briefly, each model was initialized into a pipeline object; parameter grids were defined for each to optimize based on a set of hyperparameters; tuning of the hyperparameters was performed with GridSearchCV using the r-squared (R2) to assess performance; and cross-validation of this process was performed with cross_val_score, resulting in k=5 folds for the inner grid search cross-validation, then k=10 folds for the outer cross-validation, with the mean r-squared of those five k=5 grid search cycles used to determine the best performing model.

This procedure identified the Lasso model with polynomial features as the best performing model (hyperparameters: alpha=0.1, tolerance=0.0001). The mean performance of the model during model selection as measured with the r-squared metric was 75.0±0.05. Finally, the best model selected through the above process was retrained on the full train dataset (R2=84.0). The test split was used to assess the final regression performance on out of sample control data with this model. In order to assess the mean predicted age of EP patients, the model was used to predict the ages of these subjects based on their T1/T2 ratio values. To test if there was a statistically significant difference in the mean predicted ages between the control and EP groups, an unpaired t-test was performed on the residuals of the age predictions for each group. In this case, the ages of the control group were restricted to those within the range of the ages of the EP group. The mean predicted ages of both groups were visualized as violin plots with error bars. To assess whether the thalamic nuclei of EP patients more closely resemble those of younger or older subjects, the average T1/T2 ratio values of the 44 thalamic nuclei of the EP patients were correlated with those of children in a 4-year range centered at 10 years younger than the mean EP age as well as with those of adults in a 4-year range centered at 10 years older than the mean EP age. A one sample t-test of the difference between these sets of correlations was performed, assuming the null hypothesis of a mean difference of zero. To visualize the difference as measured with Pearson correlation coefficients, a violin plot of the difference was created.

### PET neurotransmitter system maps

The neuromaps Python toolbox ^63^ was used as the source for 32 group-averaged PET maps. PET neurotransmitter system maps of receptors, transporters, and presynaptic dopamine-synthesis capacity from more than 1,200 healthy participants across 22 receptor and transporter targets in the dopamine, serotonin, glutamate, GABA, acetylcholine, opioid, cannabinoid, histamine, and norepinephrine systems were compiled ^64–89^. The full list of targets is as follows: 5HT1a/5HT1b/5HT2a/5HT4/5HT6 (serotonin receptor subtypes), D1/D2 (dopamine receptors), GABAa (γ-aminobutyric acid type A receptor), mGluR5 (metabotropic glutamate receptor 5), KOR (κ-opioid receptor), FDOPA (6-[18F]fluoro-L-DOPA [dopamine synthesis capacity]), A4B2 (α4β2 nicotinic acetylcholine receptor), M1 (M1 muscarinic acetylcholine receptor), DAT (dopamine transporter), H3 (histamine H3 receptor), VAChT (vesicular acetylcholine transporter), 5HTT (serotonin transporter), CB1 (cannabinoid receptor type 1), NMDA (N-methyl-D-aspartate receptor), GABAa5 (α5-subunit GABA-A receptor), NET (norepinephrine transporter), and MU (μ-opioid receptor). The binding potential of the radiotracer, such as radioligand tracer [11C]Flumazenil for GABA, was used as a proxy for receptor density or transporter.

Each map was resampled into the thalamic atlas space ^29^ using trilinear interpolation. Those nuclei exhibiting NaN values were excluded while those that were zero-valued were retained to avoid excluding cases of true low radioligand binding. The mean value in each nucleus was extracted and z-scored across each individual map. These mean values per nucleus were then averaged in the case of multiple source maps. The left and right CL and VAmc nuclei were excluded due to too few voxels with PET coverage upon resampling. A total of 40 nuclei survived. For molecular targets with more than one same-tracer map available (5HT1b, 5HTT, D2, GABAa, mGluR5, VAChT, MU, NET), the sample-weighted mean across the maps was calculated (see Supplementary Table 1 for specific maps used).

### Univariate analysis of neurotransmitter systems

To test whether the density of any of the 22 neurotransmitter system map measures aligned with T1/T2 ratio-defined tissue properties, Spearman rank correlation, which is robust to outliers and captures monotonic associations, was used to describe the spatial correlation between each system density and the mean developmental rate and absolute EP-NT difference of each nucleus. Permutation testing was used to assess the statistical significance of the observed correlations between the developmental rate and EP-NT difference values and the receptor/transporter density maps. Then, in addition, since smoothing was applied to both maps and thalamic nuclei exhibit spatial autocorrelation due to adjacency, BrainSMASH ^90^ was used to generate spatially constrained surrogate maps for surrogate-based spatial-null testing; briefly, it functions by generating a variogram from the empirical map, randomizing the topography of the empirical map, reintroducing spatial autocorrelation by smoothing the generated map using a distance-dependent kernel, computing a variogram from this map, regressing the new variogram on the empirical one, quantifying with sum of squared error, repeating until the value of k that minimizes SSE is determined, and finally using the best k to generate a surrogate map. To ensure a strictly positive p-value, given the possibility of producing an empirical p-value of zero under a finite number of permutations, the finite-permutation correction was used ^91^. We generated 10,000 surrogate maps. To control the false-discovery rate (FDR) across the 22 BrainSMASH-derived empirical p-values for the 22 receptor/transporter targets, the Benjamini–Hochberg (BH) procedure was applied within each analysis ^92^. To assess leave-one-nucleus-out stability, each correlation was recomputed on a leave-one-nucleus-out basis, which confirmed associations remained significant across all iterations. To test the extent of the developmental and EP neurotransmitter system profile overlap, the Spearman correlation between developmental rate and EP system profile correlations was computed (r=0.102, p>0.05). Bootstrap 95% confidence intervals (5000 resamples) were calculated for each correlation.

### Multivariate latent neurochemical axis

Considering the number of nuclei and neurotransmitter system maps, we decided to assess whether weighted combinations of systems would more strongly align with the EP-NT differences compared to single systems. Supervised decomposition using partial least squares (PLS) regression was used as our primary measure. The full matrix of 40 nuclei by 22 neurotransmitter systems was used as the input X with the EP-NT difference as y. These analyses were implemented in Python using the scikit-learn package.

PLS regression analysis was used in order to derive the latent weighted neurotransmitter system axes most strongly predictive of the tissue difference. To determine the number of PLS components k, we used LOOCV where the PLS models were fit on the 40 nuclei row x 22 receptor column input matrix, X, and used to predict the EP-NT difference vector, y, on a leave-one-nucleus-out basis. We first z-scored the input matrix and y per fold. We then regressed the EP-NT difference on the input matrix by PLS regression. The number of PLS components, k, was selected using the parsimony rule such that selected number of components was the smallest k that resulted in a LOOCV Pearson’s r that was within 0.02 of the highest Pearson’s r (k=5, r=0.73); see Fig 4b for the LOOCV-r curve). The presented Pearson coefficient is the correlation between the observed and LOOCV predicted values of the per-nucleus EP–NT difference vector. To compute the empirical permutation p-values, the nucleus labels of the EP–NT difference vector were shuffled and the full LOOCV procedure was performed for 5000 permutations, generating a null distribution of Pearson coefficients. As previously discussed, to address spatial autocorrelation BrainSMASH was used to generate a spatial null distribution based on 5000 spatial-autocorrelation-preserving surrogate maps by performing the PLS LOOCV procedure on each map. The PLS model was specified with the previously selected k to confirm that the cross-validated model performance remained significant when spatial autocorrelation was preserved.

To assess out-of-sample model performance, nested cross-validation with spatial blocking was performed. Briefly, in the outer spatially blocked fold, one left and right homotopic nucleus pair was held out resulting in 20 folds, one per each of the 20 pairs; then the remaining 38 nuclei were used in the inner LOOCV to select k as previously described; the PLS model with the inner CV-selected k specified was then trained on the 38 nuclei; and then the model was used to predict the held-out nuclei. This repeated for all 20 folds with each pair prediction concatenated to form a 40-nuclei predicted EP-NT difference vector, which was correlated with the observed EP-NT difference vector (r=0.56).

In addition to this assessment of overall model performance on predicting EP-NT difference over chance, bootstrap ratios (BSR) were computed to estimate which individual neurotransmitter system coefficients were reliable to interpret (Efron, 1979). In order to do this, the 40 nuclei were resampled with replacement (5,000 resamples) and the PLS model refit on each resample. The coefficients were sign-aligned to the original full-sample PLS model coefficients since PLS signs can flip arbitrarily, the coefficient variability estimated, and then the BSR per neurotransmitter system calculated as the original model coefficient divided by the bootstrap standard error of that distribution, which was estimated as the standard deviation of the sign-aligned bootstrap coefficients across resamples. The bootstrap ratio threshold for reliability for a given system coefficient was |BSR|>2.5 (approximately p=0.01 under a normal approximation), as a threshold indicating to what degree the magnitude of the coefficient exceeds the jitter under resampling. The BSR equation is

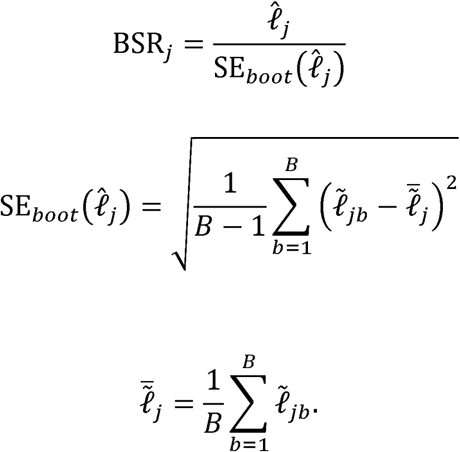

where ℓ̂*_j_* is the original full-sample PLS coefficient for system *j*, ℓ̃*_jb_* is the sign-aligned bootstrap coefficient for system *j* in bootstrap resample *b, B* = 5000, and SE*_book_* (ℓ̂*_j_*) is the bootstrap standard deviation of the sign-aligned bootstrap coefficients for that system. The PLS model with the selected number of components specified was fit on all nuclei. The PLS regression coefficients and the in-sample receptor-predicted EP-NT difference mean-centered across nuclei were extracted for later analyses. This predicted EP-NT difference is the sum of the weighted combination of all k components selected from the cross-validated prediction criterion. In the case of a k=1 model, the predicted vector is approximately equal to the single component vector multiplied by a constant weight. Single components of the multicomponent model were not interpreted individually since they are expected to exhibit high instability at low N. A given component is computed as

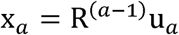

or

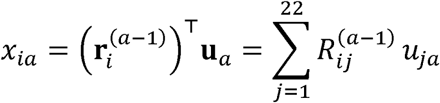

where **x***_a_* is the nucleus score vector for component *a, x_ia_* is nucleus *i*’s score on component 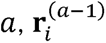 is the row *i* of the residual input matrix after components 1, …, *a* − 1 have been removed, **R**^(*a*−1)^ is the z-scored residual input matrix after previous components are removed, and **u**_a_ is the 22-receptor weight vector for component *a*. A high positive *x_ia_* means that nucleus *i*’s receptor profile aligns strongly with component *a*’s receptor pattern, so if component *a* positively weights certain receptors then that nucleus will tend to have higher values for those receptors and if component *a* negatively weights certain receptors then that nucleus will tend to have lower values for those receptors. On the other hand, a high negative *x_ia_* means that nucleus *i*’s receptor profile aligns in the opposite direction to component *a*’s receptor pattern, so if component *a* positively weights certain receptors then that nucleus will tend to have lower values for those receptors, whereas if component *a* negatively weights certain receptors then that nucleus will tend to have higher values for those receptors. The weighted sum of the components is

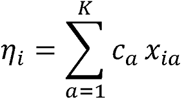

where *c_a_* is the single scalar coefficient mapping component *a*’s scores to the predicted EP-NT tissue value for that nucleus, K is the number of components, and *x_ia_* is nucleus *i*’s score on component *a*. A high positive *η_i_* means that nucleus *i*’s has a receptor profile that expresses the multicomponent PLS receptor pattern in a direction that increases the predicted EP-NT tissue difference.

The PLS-predicted EP−NT tissue difference at each nucleus is computed as

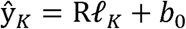

or

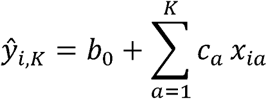

where ℓ is the vector of the 22 receptor coefficients learned by the fitted K-component PLS model and R is the z-scored 40 nuclei by 22 receptor input matrix, K is the number of components, *x_ia_* is nucleus *i*’s score on component *a, c_a_* is the single scalar coefficient mapping component *a*’s scores to the predicted EP-NT value for that nucleus, and *b*_0_ is the fitted intercept, which is approximately equal to the mean observed EP−NT tissue difference across the 40 nuclei. This value is mean-centered and interpreted as how strongly a nucleus’s receptor profile aligns with the PLS-derived multicomponent receptor pattern associated with EP-NT tissue difference.

For each subject, for each nucleus the mean tissue value within the nucleus was z-scored across subjects and weighted by the nucleus’s PLS predicted nucleus value, then all weighted nuclei values were summed, resulting in a single neurotransmitter system-weighted value (receptor score) for each subject. The formula for this value is

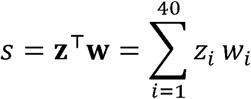

where *s* is a subject’s scalar score, *z* is their 40-value mean nucleus tissue value vector, *w* is a their 40-value mean-centered PLS-predicted EP−NT tissue difference, *z_i_* is their mean tissue value at nucleus *i*, and *w_i_* is their PLS-predicted EP−NT tissue difference at nucleus *i*. This receptor score is interpreted as how strongly a subject’s overall thalamic tissue pattern aligns with the latent neurochemical profile.

### Psychotic symptom analyses

Psychiatric symptoms were measured using the Positive and Negative Syndrome Scale (including positive, negative, and general subscales), Clinical Assessment Interview for Negative Symptoms, Montgomery–Åsberg Depression Rating Scale, and Young Mania Rating Scale ^93–96^. In the primary analysis in the EP group to assess how receptor score, which quantifies the degree to which a patient’s thalamic tissue aligns with the latent neurochemical profile, predicts psychotic symptoms, we fit 7 separate OLS regression models (HC3 robust standard errors) with receptor score as the independent variable, antipsychotic chlorpromazine-equivalent dose (CPZ) as a covariate, and each of the 7 symptom scales as dependent variables. All predictors were standardized to allow for comparison. For a specificity control to confirm that the relationship was attributable to receptor weighting rather than to overall tissue, another set of 7 models was fit with the mean of the z-scored tissue across nuclei as the independent variable, CPZ as a covariate, and the 7 symptom scales as dependent variables. The primary analysis was BH-FDR-corrected across all 7 tests.

We computed an EP risk score as a measure of how much a given subject’s thalamic tissue pattern aligns with the direction of the NT-EP tissue difference map. For each subject, the mean tissue value within each nucleus was z-scored relative to the NT subjects and the remaining EP subjects, weighted by the nucleus’s signed NT-EP tissue difference value, then all weighted nucleus values were summed. A higher score indicates that the subject’s tissue pattern is more aligned with the signed NT-EP tissue-difference pattern: relatively higher tissue in nuclei where EP < NT and relatively lower tissue in nuclei where EP > NT.

## Data and Code Availability

The HCP data are available from the Connectome Coordination Facility (https://www.humanconnectome.org/). The mrGrad software used to quantify and create visualizations of thalamic gradients is available at the Mezer Lab Github (https://github.com/MezerLab/mrGrad). Other visualization code is available at the author’s Github (https://github.com/o-sing/papers).

**Supplementary Figure 1:**
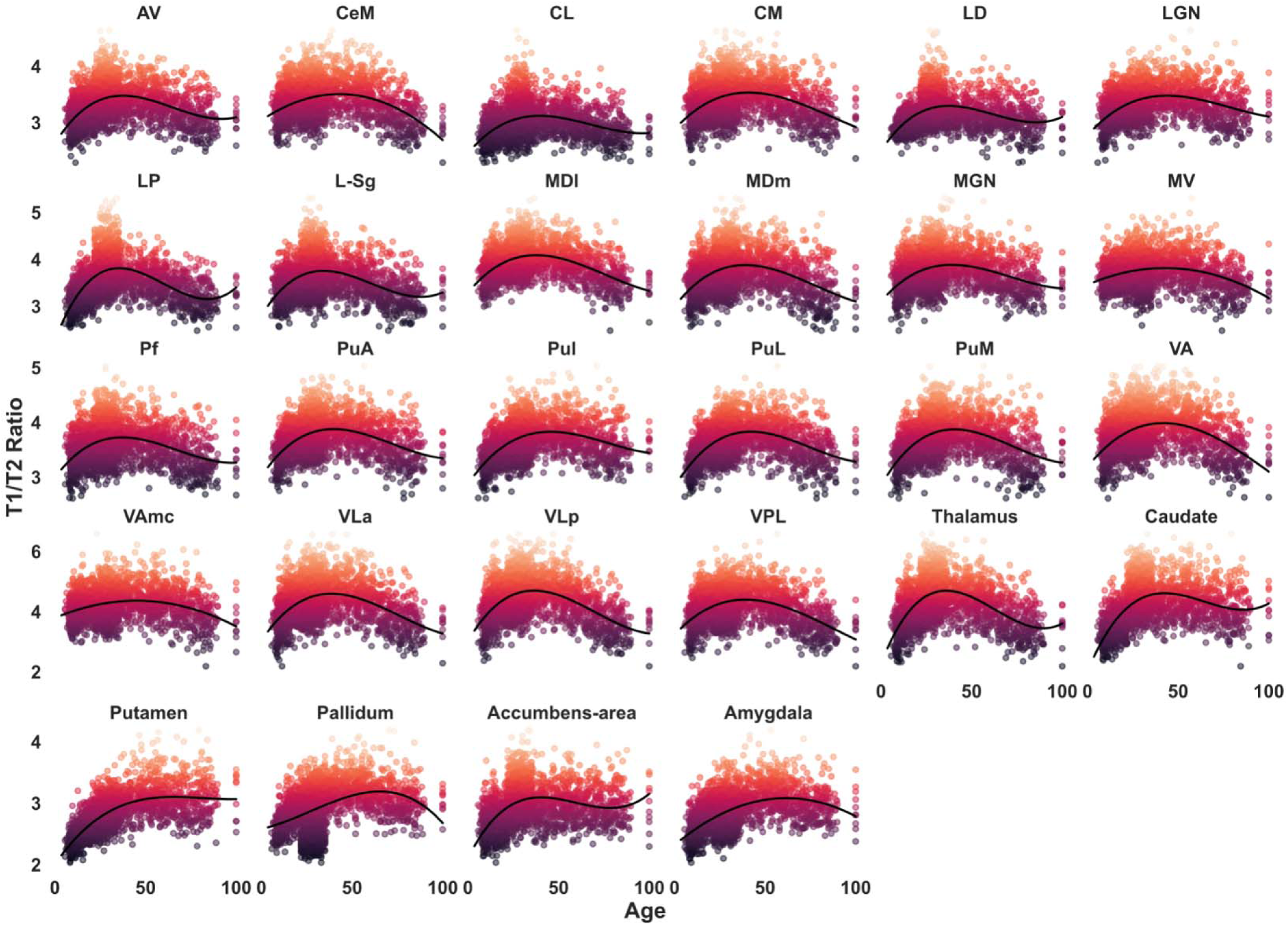
Developmental trajectories of all left hemisphere subcortical ROIs. Scatterplots of T1/T2 ratio as a function of age in all subcortical ROIs. Each dot represents the mean T1/T2 ratio value in a given participant’s nucleus or other subcortical structure. Black line denotes best-fit polynomial across participants.

**Supplementary Figure 2:**
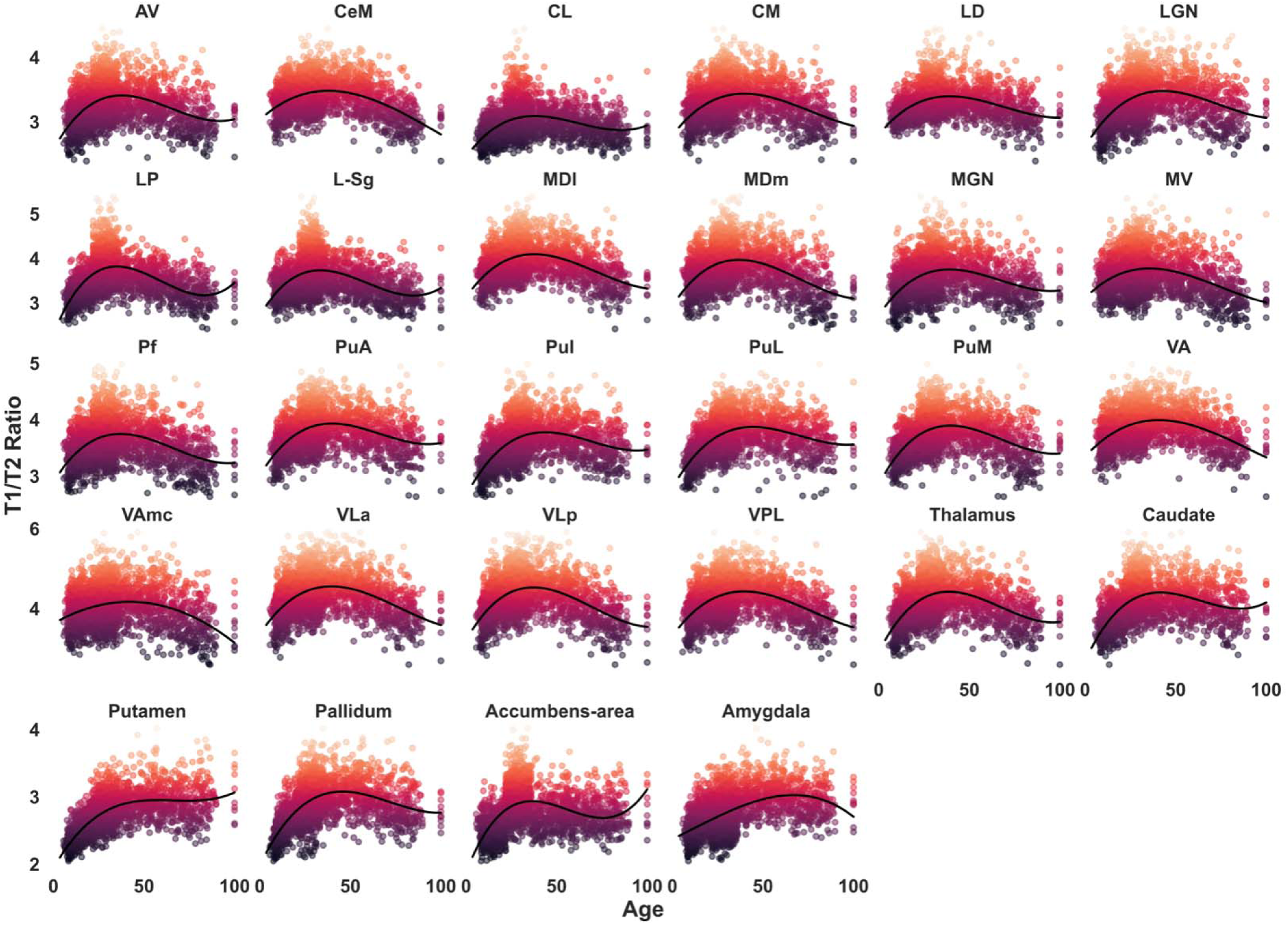
Developmental trajectories of all right hemisphere subcortical ROIs. Scatterplots of T1/T2 ratio as a function of age in all subcortical ROIs. Each dot represents the mean T1/T2 ratio value in a given participant’s nucleus or other subcortical structure. Black line denotes best-fit polynomial across participants.

**Supplementary Table 1.**
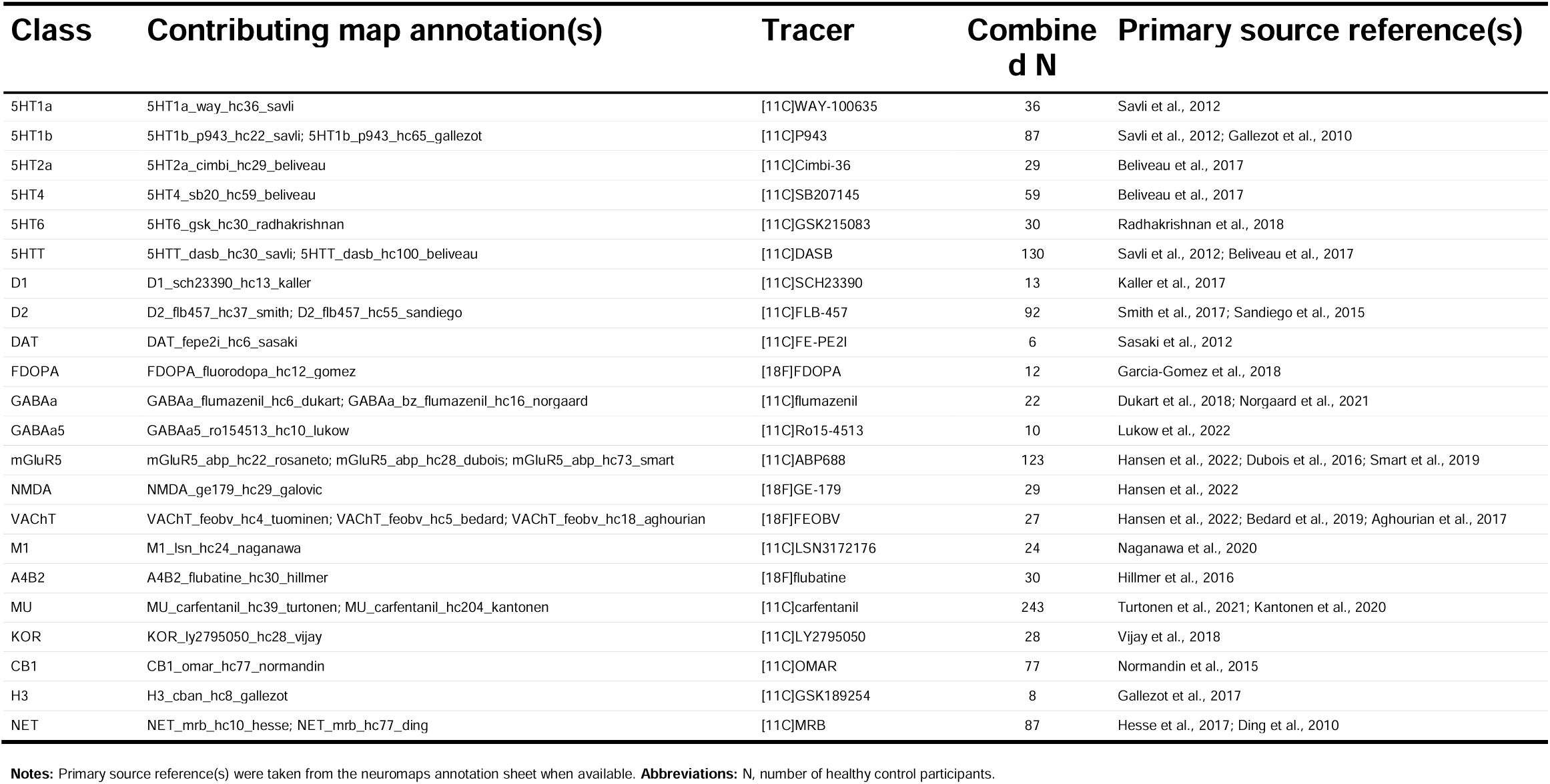
Neurotransmitter receptor and transporter panel by receptor class For receptor classes represented by more than one map of the same radiotracer, maps were combined by sample-size-weighted averaging of z-scored values. Combined N is the sum of healthy control participants across the contributing source maps.

**Supplementary Table 2.**
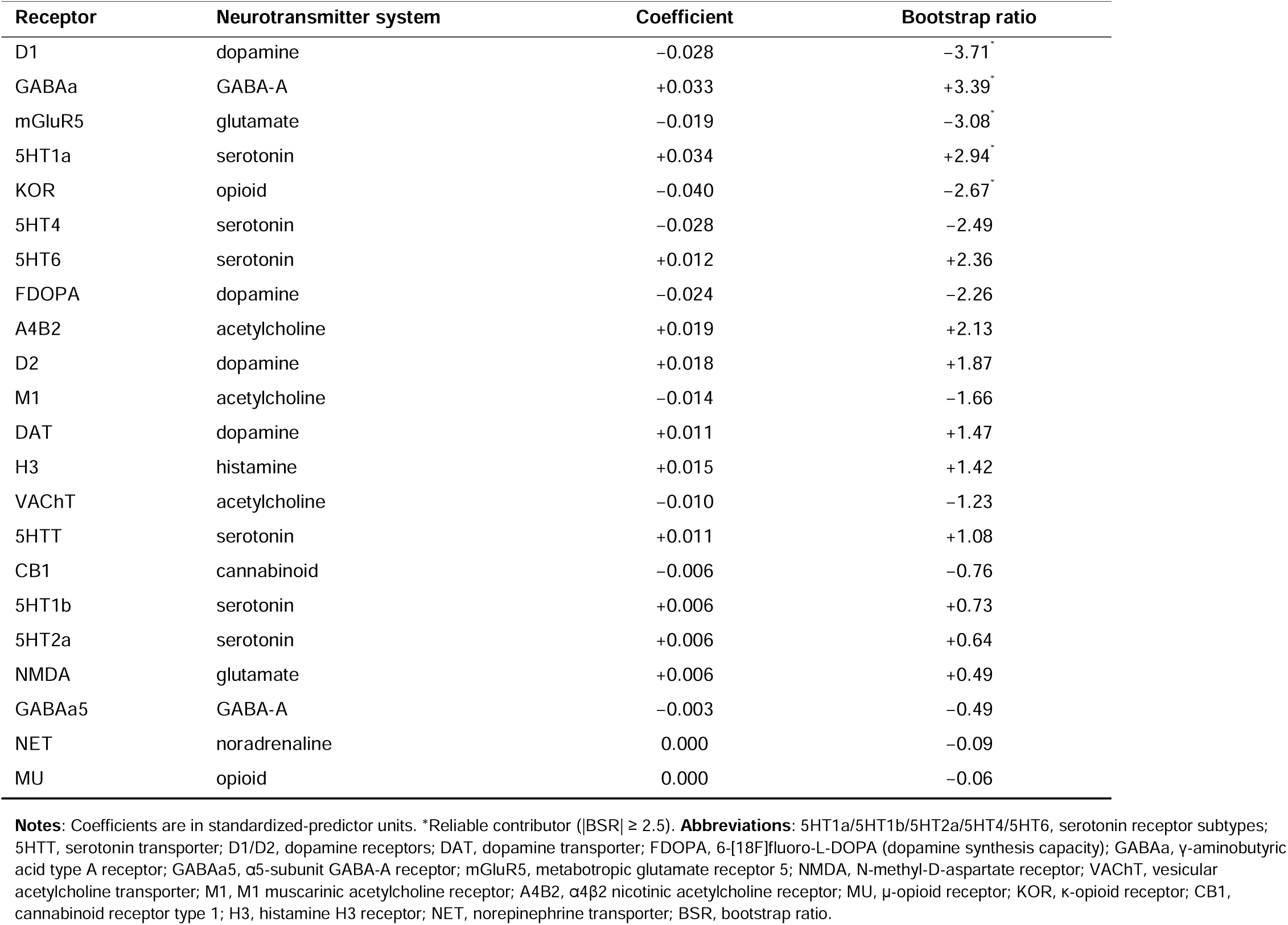
Reliability of receptor and transporter contributions to the latent neurochemical profile.

